# Activity-dependent regulation of the proapoptotic BH3-only gene *egl-1* in a living neuron pair in *C. elegans*

**DOI:** 10.1101/548925

**Authors:** Jesse Cohn, Vivek Dwivedi, Nicole Zarate, H Robert Horvitz, Jonathan T Pierce

## Abstract

The BH3-only family of proteins is key for initiating apoptosis in a variety of contexts, and may also contribute to non-apoptotic cellular processes. Historically, the nematode *Caenorhabditis elegans* has provided a powerful system for studying and identifying conserved regulators of BH3-only proteins. In *C. elegans*, the BH3-only protein EGL-1 is expressed during development to cell-autonomously trigger most developmental cell deaths. Here we provide evidence that *egl-1* is also transcribed after development in the sensory neuron pair URX without inducing apoptosis. We used genetic screening and epistasis analysis to determine that its transcription is regulated in URX by neuronal activity and/or in parallel by orthologs of Protein Kinase G and the Salt-Inducible Kinase family. Because several BH3-only family proteins are also expressed in the adult nervous system of mammals, we suggest that studying *egl-1* expression in URX may shed light on mechanisms that regulate conserved family members in higher organisms.

## INTRODUCTION

Members of the BH3-only family of proteins are essential regulators of cell death in nearly all metazoans. Each member of this family shares the “<u>B</u>CL-2 <u>h</u>omology” domain BH3, which allows them to bind the anti-apoptotic members of the BCL-2 family and block their activity, thereby initiating apoptosis (LETTRE AND HENGARTNER 2006; GIAM *et al.* 2008). In mammals, there are at least eight BH3-only proteins, with some estimates ranging higher (AOUACHERIA *et al.* 2005; HAPPO *et al.* 2012). These proteins are regulated in a number of ways, such as transcriptionally (ODA *et al.* 2000; NAKANO AND VOUSDEN 2001; STURM *et al.* 2006; BORENSZTAJN *et al.* 2007), post-transcriptionally (VENTURA *et al.* 2008; SHERRARD *et al.* 2017), and post-translationally (VERMA *et al.* 2001; LOWMAN *et al.* 2010). This complex regulation allows for cells to properly induce cell death during development and in response to a wide variety of contexts and circumstances. Additionally, mounting evidence suggests that some BH3-only proteins may contribute to non-apoptotic cellular processes as well, such as glucose metabolism (DANIAL *et al.* 2003; DANIAL *et al.* 2008; GIMENEZ-CASSINA *et al.* 2014), autophagy (MAIURI *et al.* 2007; SINHA *et al.* 2008; LINDQVIST *et al.* 2014), and lipid transport (ESPOSTI *et al.* 2001). Despite their important role in initiating programmed cell death and likely other cellular processes, our understanding of the factors that regulate BH3-only proteins is incomplete.

The nematode *C. elegans* has historically played an important role in understanding the mechanisms of programmed cell death and the role of BH3-only proteins (LETTRE AND HENGARTNER 2006). Genetic screens in the 1980s and 1990s using *C. elegans* uncovered a core pathway through which programmed cell death is initiated and executed (ELLIS AND HORVITZ 1986; YUAN AND HORVITZ 1990; HENGARTNER *et al.* 1992; YUAN AND HORVITZ 1992; CONRADT AND HORVITZ 1998), the main features of which are well conserved all the way through mammals (CZABOTAR *et al.* 2014). The primary initiator of this cell death pathway was found to be a BH3-only protein called EGL-1 (<u>eg</u>g-laying defective-<u>1</u>). A gain-of-function mutation in the regulatory region of *egl-1* causes the inappropriate death of neurons in the egg-laying circuit, while subsequent loss-of-function mutations in *egl-1* prevented this cell death (CONRADT AND HORVITZ 1998; CONRADT AND HORVITZ 1999). EGL-1 was found to be necessary for most developmental cell deaths in the worm, and sufficient to induce apoptosis cell-autonomously when expressed ectopically *in vivo* (CONRADT AND HORVITZ 1998; CHANG *et al.* 2006; LOMONOSOVA AND CHINNADURAI 2008). So far, evidence exists for *egl-1* regulation at the transcriptional and the post-transcriptional level (NEHME AND CONRADT 2008; NEHME *et al.* 2010; SHERRARD *et al.* 2017). By focusing on individual cells that die during development, some of the transcription factors and pathways that regulate *egl-1* transcription in apoptotic cells have been found (ELLIS AND HORVITZ 1991; METZSTEIN *et al.* 1996; CONRADT AND HORVITZ 1999; THELLMANN *et al.* 2003; LIU *et al.* 2006; POTTS *et al.* 2009; HIROSE *et al.* 2010; WINN *et al.* 2011; HIROSE AND HORVITZ 2013; SHERRARD *et al.* 2017). Notably, homologs of nearly all of these factors have also been implicated in apoptosis in higher animals, suggesting that studying *egl-1* regulation in *C. elegans* is a fruitful approach for identifying mechanisms important in cell death pathways in other systems (WALLIS *et al.* 1999; XU *et al.* 1999; GINSBERG 2002; RUIZ I ALTABA *et al.* 2002; WU *et al.* 2005; DENIAUD *et al.* 2006; WONG *et al.* 2007; SITWALA *et al.* 2008).

Here we describe the first non-apoptotic example of endogenous *egl-1* transcription in living neurons in *C. elegans* after development. We find that *egl-1* mRNA is present in a very small subset of neurons in *C. elegans* and persists into adulthood – most consistently and reliably in the neuron pair URX, the main oxygen sensing neurons in *C. elegans*. The URX neurons are necessary for worms to orient properly to different oxygen levels in their environment (GRAY *et al.* 2004; ZIMMER *et al.* 2009; BUSCH *et al.* 2012), and are also involved in other various aspects of worm physiology such as fat homeostasis and immune responses (STYER *et al.* 2008; WITHAM *et al.* 2016). URX senses oxygen via a heterodimer of soluble guanylyl-cyclases, GCY-35 and GCY-36, which produces cGMP in response to binding oxygen. This cGMP then gates calcium currents via the cyclic-nucleotide gated ion channels CNG-1 and TAX-4 commensurate with the level of environmental oxygen (GRAY *et al.* 2004; BUSCH *et al.* 2012; COUTO *et al.* 2013). With the use of a fluorescent transcriptional reporter for *egl-1*, we find that sensory transduction in URX is necessary for *egl-1* transcription. This transcription can also be regulated in parallel by two protein kinases: EGL-4, the worm homolog of Protein Kinase G, and KIN-29, the worm Salt-Inducible Kinase family homolog, as well as by at least one more unknown pathway.

Because the study of *egl-1* regulation has historically yielded insight into factors that control apoptosis in higher animals, we propose that this novel adult transcription of *egl-1* in URX may further our understanding of roles and regulatory mechanisms of BH3-only proteins in apoptotic and potentially non-apoptotic contexts.

## MATERIALS AND METHODS

### Strains Used

The following strains were used: N2 Bristol as wild type; JPS600 *vxEx600[Pegl-1::mCherry Punc-122::GFP]*; JPS601 *vxIs601[Pegl-1::mCherry Punc-122::GFP]*; JPS602 *vxEx602[Pegl-1::mCherry Punc-122::GFP Pgcy-32::GFP]*; JPS620 *ced-6(n1813) III; vxEx602[Pegl-1::mCherry Punc-122::GFP Pgcy-32::GFP]*; MT22516 *him-8(e1489) V; nIs343[Pegl-1::4xNLS::GFP]*; MT8735 *egl-1(n1084n3082) V*; RB1305 *egl-1(ok1418) V*; CX4148 *npr-1(ky13) X*; JPS1101 *egl-1(n1084n3082) V; npr-1(ky13) X*; JPS879 *vxEx879[Pgcy-32::GFP Punc-122::GFP]*; JPS1106 *egl-1(n1084n3082) V; vxEx1106[Pgcy-32::GFP Pfat-7::GFP]*; JPS1031 *vxEx1031[Pgcy-32::RAB-3::GFP Pgcy-32::mCherry]*; JPS1070 *egl-1(n1084n3082) V; vxEx1031[Pgcy-32::RAB-3::GFP Pgcy-32::mCherry]*; JPS696 *cng-1(vx3) V; vxIs601[Pegl-1::mCherry Punc-122::GFP]*; JPS803 *cng-1(vx3) V; vxIs601[Pegl-1::mCherry Punc-122::GFP]*; *vxEx803[Pgcy-32::CNG-1(+) Peft-2::GFP]*; JPS793 *gcy-35(ok769) I; vxEx600[Pegl-1::mCherry Punc-122::GFP]*; JPS909 *gcy-36(db42) X; vxEx600[Pegl-1::mCherry Punc-122::GFP]*; JPS812 *egl-19(n582) IV; vxEx600[Pegl-1::mCherry Punc-122::GFP]*; JPS839 *egl-19(n2368) IV; vxEx600[Pegl-1::mCherry Punc-122::GFP]*; JPS937 *egl-4(n477) IV; vxIs601[Pegl-1::mCherry Punc-122::GFP]*; JPS1124 *kin-29(oy38) X; vxIs601[Pegl-1::mCherry Punc-122::GFP]*; JPS1040 *egl-4(n477) IV; kin-29(oy38) X; vxIs601[Pegl-1::mCherry Punc-122::GFP]*; JPS1020 *egl-4(n477) IV; muIs102[Pgcy-32::GFP] V*; JPS912 *mef-2(gv1) I; vxIs601[Pegl-1::mCherry Punc-122::GFP]*; JPS893 *egl-4(vx19) IV; cng-1(vx3) V; vxIs601[Pegl-1::mCherry Punc-122::GFP]*; JPS1127 *cng-1(jh111) V; kin-29(oy38) X; vxIs601[Pegl-1::mCherry Punc-122::GFP]*; JPS1126 *gcy-35(ok769) I; egl-4(n477); kin-29(oy38) X; vxIs601[Pegl-1::mCherry Punc-122::GFP]*

### Molecular Biology and Transgenic Strain Construction

The *egl-1* reporter construct was generated by long PCR fusion of three fragments: 1042-bp upstream of the *egl-1* start codon amplified from N2 DNA lysate, worm-optimized mCherry amplified from plasmid pCFJ90, and 5744-bp of the 3’ downstream region of *egl-1* genomic DNA including the 3’UTR of *egl-1* amplified from N2 lysate (SHEVCHUK *et al.* 2004). Primers used to generate the final fused product were: AGGCCTGATCATAGTTTCTGCCATTT and ATCCCTAACATATTTCTCAAAGATACAAATGTCATC.

To generate the strain JPS601 with the integrated *egl-1* reporter transgene *vxIs601*, we first transformed the above PCR product into N2 WT by microinjection at a concentration of 2 ng/µl along with a *Punc-122::GFP* co-injection marker to generate strain JPS600 carrying the extra-chromosomal array *vxEx600*. This array was then integrated by UV irradiation using a UV Stratalinker 2400 (Stratagene), and the resulting integrated strain was then outcrossed to N2 six times before any further use (MARIOL *et al.* 2013).

URX neuron labeling constructs were generated by using an 876-bp fragment of the *gcy-32* promoter established from earlier studies (YU *et al.* 1997). The promoter fragment was amplified from N2 gDNA and then PCR fused with either worm-optimized mCherry amplified from pCFJ90 or worm-optimized GFP amplified from pPD95.75. The fused products were then subcloned into the pCR-Blunt vector using a Zero Blunt PCR Cloning Kit from ThermoFisher Scientific. These constructs were used in injection mixes at a concentration of 20 ng/µl.

The URX-targeted synaptic marker RAB-3::GFP was generated by digesting plasmid NM1028 with *Pst*I and *Nco*I and inserting the *gcy-32* promoter with Gibson Assembly. This construct was injected at a concentration of 25 ng/µl. NM1028 was a kind gift of Michael Nonet (Washington University in St. Louis).

The construct to cell-specifically rescue *cng-1* in URX was generated by amplifying the full coding region of *cng-1* and the *gcy-32* promoter each from N2 gDNA. These were then assembled by Gibson Assembly with a pPD95.75 backbone that had been digested with *Xba*I and *Eco*RI. This was then injected at a concentration of 20 ng/µl.

Mutations were followed in crosses by PCR genotyping and/or by phenotype where applicable.

### Mutagenesis and Mutant Identification

To screen for mutations that affect expression of *vxIs601* in URX, we mutagenized the otherwise wild-type strain JPS601 with 0.5mM N-ethyl-N-nitrosourea (ENU) in M9 buffer for 4 hours and then examined F2 progeny using a fluorescence dissection microscope. We looked for animals with a loss or diminishment of mCherry expression in URX. We followed the strategy outlined in Zuryn, et. al. (2010), to identify candidate causal mutations. We identified allele *vx3* as an A-to-C mutation 651 bp from the start codon of *cng-1* predicted to convert the aspartic acid at position 162 to an alanine.

For our reversion screen, *cng-1(vx3)* mutant animals were backcrossed to strain JPS601 six times and then subjected to ENU treatment as above. F2 progeny were screened on a fluorescence dissection microscope to look for recovery of fluorescence in URX. All revertant mutants displayed a slight egg-laying defect and a longer body than wild-type – common characteristics of the *egl-4* mutant. Revertant mutants failed to complement the *egl-4(n477)* allele when tested for *vxIs601* expression in URX.

### General Microscopy

Worms were mounted on 2% agarose pads and anesthetized with 30-mM sodium azide in NGM. Epifluorescent images were taken on an Olympus IX51 inverted microscope equipped with an X-Cite FIRE LED Illuminator (Excelitas Technologies Corp.) using an Olympus UPlanFL N 40X/0.75 NA objective and QCapture Pro 6.0 software. Confocal images were taken with a Zeiss LSM 710 microscope equipped with a Plan-Apochromat 40x/1.4 Oil objective and Zen Software.

### *egl-1* Reporter Fluorescence Quantification and Oxygen Experiments

For fluorescence quantification of the *egl-1* reporter in URX, animals were synchronized by timed egg laying and then imaged three days later. *kin-29* mutants develop significantly slower than wild-type worms or *egl-4* mutants, however because reporter expression level is a function of the amount of time an animal is exposed to oxygen as opposed to a particular age, we compared day 1 wild-type and *egl-4* adults to L2-L3 *kin-29* animals that had been synchronized by egg-laying.

Fluorescence intensity was quantified in FIJI software as has been previously described (MCCLOY *et al.* 2014). Briefly, the cell body was outlined and then the background was subtracted to give a final fluorescence intensity for each cell. These values were then normalized to wild-type animals grown in parallel at 21% oxygen for comparison purposes. Pictures shown in figures are to show presence/absence of reporter expression; non-saturated photos were used for quantification.

For experiments in which the oxygen environment was different than 21%, worms were grown or maintained in a Modular Incubator Chamber (Billups-Rothenberg) attached to oxygen tanks containing 100% medical grade O2 (Praxair), or either 1% or 10% O2 balanced with nitrogen (Airgas).

### Single Molecule FISH

smFISH experiments were performed as described elsewhere (RAJ *et al.* 2008). Fixed mixed stage animals were incubated overnight at 30° with previously designed *egl-1* probes (JOHNSEN AND HORVITZ 2016). Animals were subsequently mounted for imaging after two washes lasting 30 minutes each. Image acquisition was performed on a Nikon TE-2000 inverted microscope with a 100x objective (Nikon, NA 1.4). A Pixis 1024 camera (Princeton Instruments) controlled by MetaMorph software (Molecular Devices) was used to detect smFISH signal and acquire images with an exposure time of 2 seconds. Images were processed and prepared for publication using ImageJ software (NIH).

### Lifespan Assays on *Pseudomonas aeruginosa*

Assays were performed as previously described (STYER *et al.* 2008). *P. aeruginosa* was grown overnight at 37° and then seeded onto NGM-agar plates modified to contain 0.35% peptone. After seeding, plates were placed at 37° overnight and moved in the morning to 25° to equilibrate temperature. At least 10 gravid adults were placed on each plate with 4-5 replicate plates per strain per assay. Worms were assessed every 4-8 hours for survival. Animals that failed to respond to touch were counted as dead, and animals that crawled up the sides of the plate and desiccated were censored. Worms were maintained at 25° and moved to fresh plates every other day over the course of the assay.

### Bordering Assays

Assays were performed as previously described (GROSS *et al.* 2014). NGM agar plates were seeded with 200µl of fresh overnight cultures of OP50 two days prior to testing. Worms were age-matched by selecting L4 animals the day before the assay. The day of the assay, 40 day 1 adults were picked to each assay plate, with at least 4-5 replicates per strain per assay. The number of animals on the lawn border was quantified one hour after addition of worms to the plate.

### Dendrite Scoring Assays

URX dendrites in day 4 adults carrying a *Pgcy-32::GFP* transgene were visualized and scored for tip morphology. Dendritic endings with a secondary branch that extended at least 5 microns from the primary branch were scored as “complex”, otherwise they were scored as “simple”. About 17% of URX dendrites in the *egl-1* mutant failed to extend all the way to the tip of the nose, possibly because of extra undead cells interfering with dendritic attachments to glia. These animals were not included in the determination of complex/simple ratios.

### Data Availability

Strains and plasmids are available upon request. The authors affirm that all data necessary for confirming the conclusions of the article are present within the article and figures.

## RESULTS

### *egl-1* is transcribed in URX neurons post-developmentally

To investigate the expression pattern of *egl-1*, we generated a transcriptional reporter construct consisting of 1.1 kb of the upstream promoter region, as well as 5.5 kb of the region downstream of the gene. The coding region of *egl-1* was replaced with a *C. elegans*-optimized version of mCherry (Figure 1A). We integrated this construct to create the transgenic reporter *vxIs601* (MARIOL *et al.* 2013). To ensure that *vxIs601* faithfully reported true *egl-1* expression, we visualized apoptotic cells in a *ced-6(n1813)* mutant background in which dead cell corpses persist due to a defect in engulfment (Figure 1A’) (LIU AND HENGARTNER 1998). We found that apoptotic cell corpses reliably showed high mCherry expression, indicating that *vxIs601* correctly reflects *egl-1* transcription in at least the developmental cell deaths we examined. For simplicity and clarity, we refer to *vxIs601* as the *egl-1* reporter from hereon.

**Figure 1.**
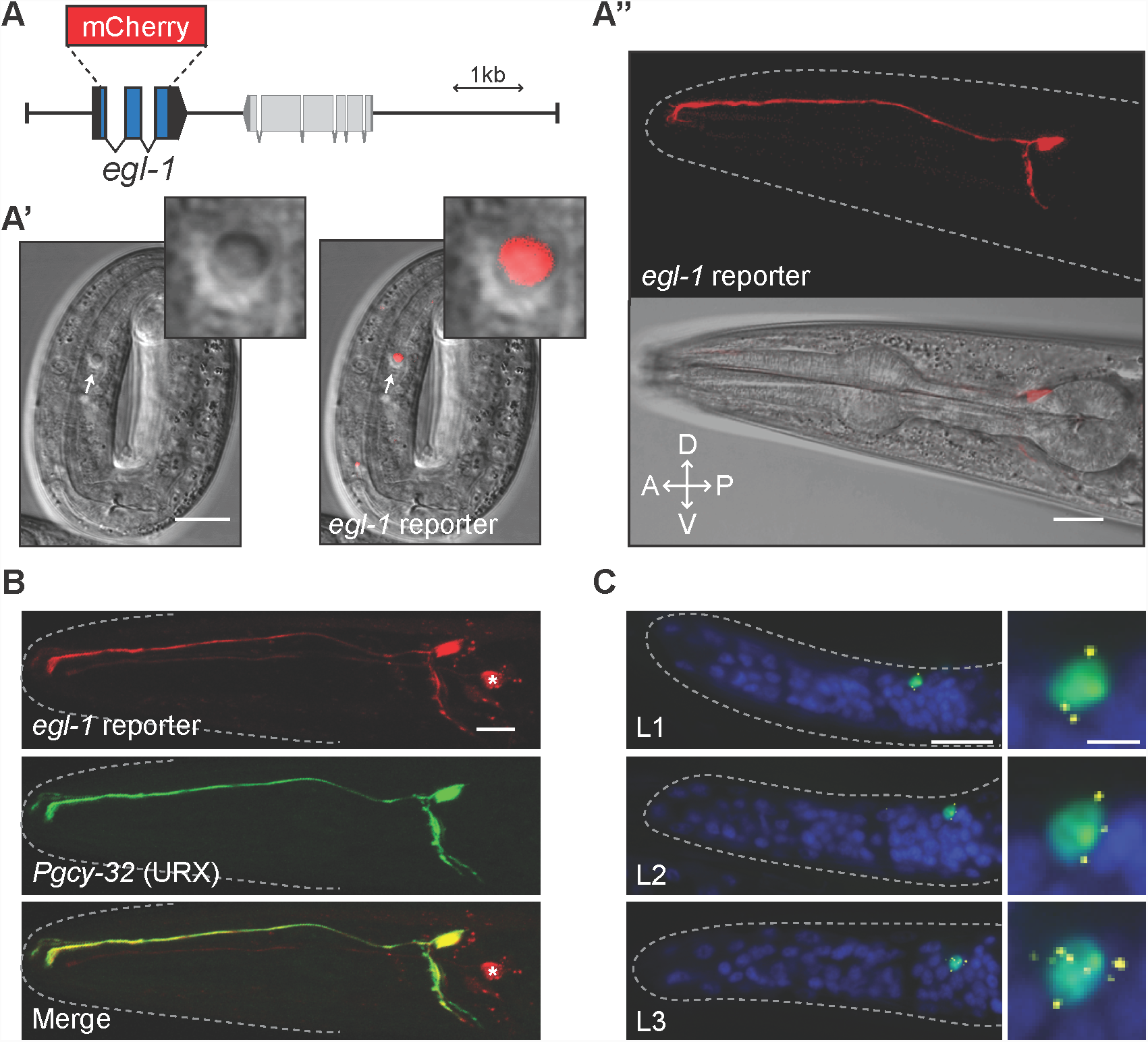
The BH3-only gene *egl-1* is transcribed post-developmentally in the URX neuron pair. **A.** Schematic representation of the *egl-1* reporter construct in which the *egl-1* coding sequence is replaced with *mCherry*. **A’.** Expression of the *egl-1* reporter construct in a *ced-6(n1813)* embryo in which dead cell corpses persist. Arrows and inset show an apoptotic cell corpse that expresses the *egl-1* reporter. **A’’.** Example of *egl-1* reporter expression in a day 2 adult wild-type worm. **B.** Co-localization of the *egl-1* reporter with a *Pgcy-32::GFP* reporter that labels the URX neuron pair. Asterisk indicates expression of the *egl-1* reporter in cells other than URX that occasionally occurs. **C.** Single molecule fluorescent *in situ* hybridization (smFISH) showing *egl-1* transcripts in animals carrying the *nIs343* transgene (*Pegl-1::4xNLS::GFP*). Blue is DAPI, *egl-1* mRNA transcripts are labeled yellow, green is GFP. Larval age of each animal is shown in white text. Scale bars in all pictures are 10 µm, except for the enlargements in C in which the scale bar is 2 µm.

To our surprise, we also found that the *egl-1* reporter was expressed in adult animals in a very small subset of neurons, and in one pair especially strongly in all animals (Figure 1A’’). This expression persisted throughout the entire life of the worm. We used cell-specific reporters and morphological characterization to identify the neurons most brightly expressing the reporter as the bilaterally symmetric neuron pair URX (Figure 1B) (YU *et al.* 1997). We also found that AQR, PQR and one of the AWCon/off neuron pair expressed the reporter less often (see Figure 1B, top, for an example). Of these neurons, the expression in the URX neuron pair was the most consistent and robust, so we chose to characterize this adult expression of *egl-1* by focusing our attention specifically on URX.

Reporter genes occasionally express in cells that do not normally express the endogenous gene because the reporter lacks regulatory regions that repress the endogenous gene (TURSUN *et al.* 2009). To test this possibility for our *egl-1* reporter, we used single molecule fluorescence *in-situ* hybridization (smFISH) to confirm whether endogenous *egl-1* mRNA transcripts could be found in URX (JOHNSEN AND HORVITZ 2016). To test the co-localization of the *egl-1* fluorescent reporter and endogenous *egl-1* mRNA, we used animals that carry the *egl-1* reporter transgene *nIs343*, which expresses nuclear-localized GFP using 6.5 kb of the upstream promoter region of *egl-1* (HIROSE AND HORVITZ 2013). These animals showed the same expression pattern as our *egl-1* reporter described above (*vxIs601*), including expression of GFP in a small number of neurons persisting past development, and especially strongly in URX. We found that the GFP expression always (8/8 worms) co-localized with the smFISH signal from probes targeted to the *egl-1* transcript (Figure 1C). Notably, this expression continued past the L2 larval stage, which is when the last somatic cell deaths in the hermaphrodite occur (SULSTON AND HORVITZ 1977).

Past studies have identified genomic regulatory regions around *egl-1* that control expression in certain apoptotic cells, so we performed promoter truncation experiments to determine whether *egl-1* expression in URX is dependent on regions previously identified as important for *egl-1* transcription in other cells (BOULIN *et al.* 2006). We identified a 20-bp region ∼85bp upstream from the *egl-1* transcriptional start site that was necessary for reporter expression in URX (Figure S1). This region has not been identified as necessary for any cell deaths examined in the worm thus far, and it is near but distinct from a previously identified binding site of the transcription factor SPTF-3 (HIROSE AND HORVITZ 2013). This suggests that *egl-1* transcription is regulated in URX by different regulatory regions than have been previously described in developmental cell deaths.

We tried several approaches to determine whether the EGL-1 protein was also produced in URX; however, due to technical difficulties we were unable to make this assessment (see Discussion for more details). Nevertheless, we conclude that endogenous *egl-1* is transcribed post-developmentally in the URX neuron pair without initiating apoptosis.

### No obvious role for *egl-1* in URX neurons

The URX neurons do not die under normal conditions. This leaves open the possibility that *egl-1* may play a non-apoptotic role in URX. To address this hypothesis, we examined the *egl-1* loss-function mutant with regard to a variety of URX- and *egl-1*-related phenotypes.

URX has an established role in the *C. elegans* innate immune response. Transgenic worms lacking the URX, AQR, and PQR neurons survive longer than wild-type animals on the pathological bacteria *Pseudomonas aeruginosa* (STYER *et al.* 2008). We found that the *egl-1* mutant has similar survival rates as wild-type on *P. aeruginosa*, indicating that *egl-1* is not necessary for URX to contribute to the worm innate immune response (Figure 2A). We also examined URX morphology to look for any signs of neurodegeneration after incubation for 24 hours on *P. aeruginosa*, but found no obvious defects in any (40/40) worms.

**Figure 2.**
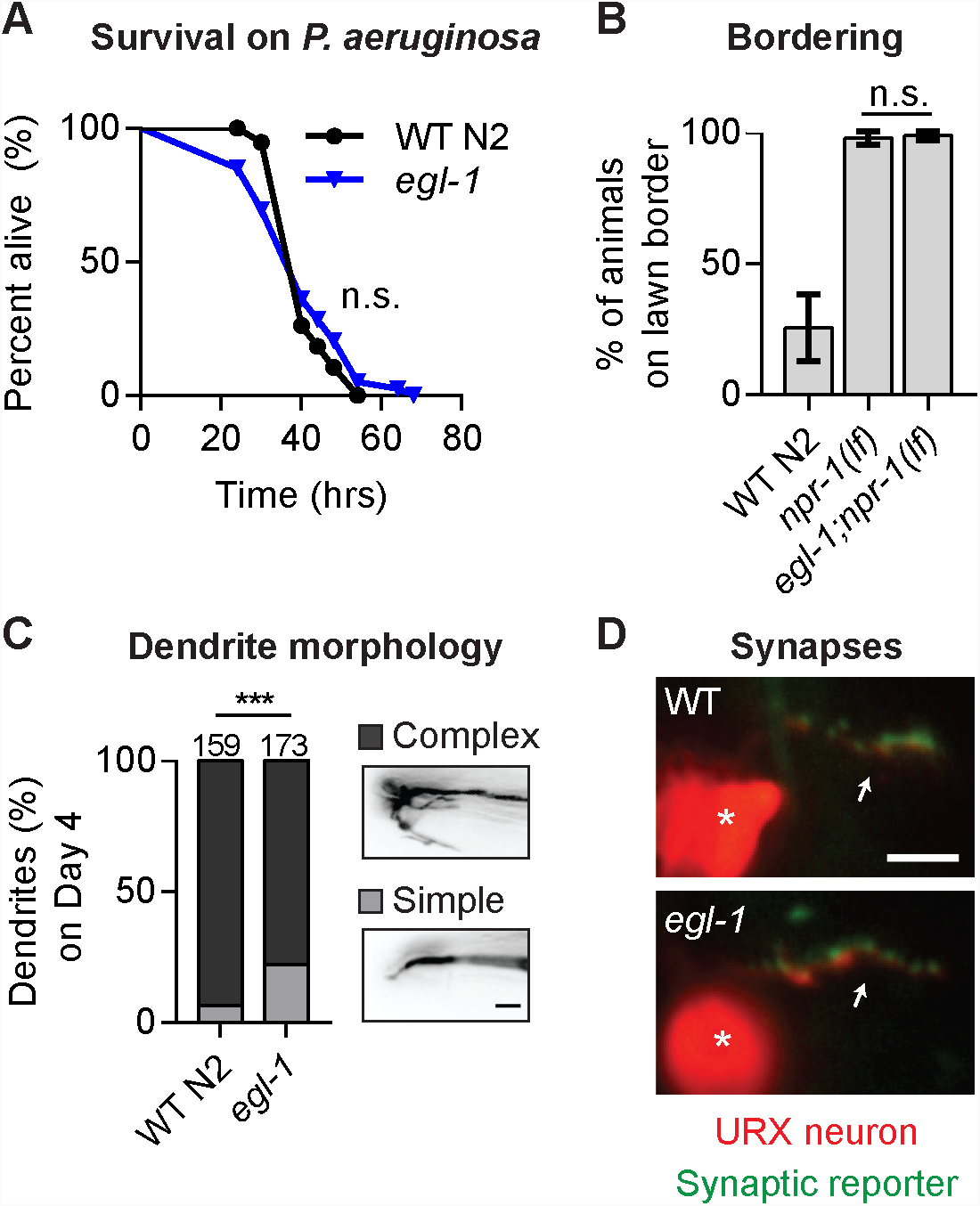
Analysis of URX- and *egl-1*-related phenotypes in wild-type and *egl-1* mutant worms. **A.** Lifespan of worms transferred as day 1 adults to pathogenic *Pseudomonas aeruginosa*. Wild type (n = 38) vs. *egl-1(ok1418)* (n = 39), p = 0.93 as determined by the Mantel-Cox test. **B.** Bordering assay of day 1 adult worms in 21% oxygen. Mean ± 95% C.I. *npr-1(ky13)* (n = 4 plates of 40 worms) vs. *egl-1(n1084n3082);npr-1(ky13)* (n = 4 plates of 40 worms), p = 0.40 as determined by Student’s t-test. **C.** Dendritic morphology assay of day 4 adults. Complex dendrites have at least one secondary branch that extends at least 5 µm from the primary dendritic stalk, simple dendrites do not. Wild-type (n = 159) vs. *egl-1(n1084n3082)* (n = 173), p < 0.01 as determined by Fisher’s exact test. **D.** Representative photos of URX synapses (green) in wild type and *egl-1(n1084n3082)* day 1 adults. Both strains carry a *[Pgcy-32::mCherry Pgcy-32::RAB-3::GFP]* reporter transgene. Arrows point to the URX axon (red) in the nerve ring, asterisks label the URX cell body. All scale bars are 5 µm.

As the main oxygen-sensing neuron in *C. elegans*, URX is necessary for the bordering behavior exhibited by *npr-1* loss-of-function mutant animals (CHEUNG *et al.* 2004; GRAY *et al.* 2004). This behavior reflects a strong aversion to surface-level oxygen (21% O2), and *npr-1* mutants lacking URX or components of the oxygen-sensing machinery in URX show a defect in bordering. However, we found that the *npr-1;egl-1* double mutant borders with the same incidence as the *npr-1* single mutant. This suggests that *egl-1* is not necessary for URX to sense oxygen in order to drive the bordering behavior (Figure 2B).

URX has been described as having a characteristic variable, branched dendritic ending at the tip of the nose (“complex” endings) (WARD *et al.* 1975; WHITE *et al.* 1986; DOROQUEZ *et al.* 2014). In our parallel submitted work, we show that URX dendritic tips fail to branch in certain genetic and environmental conditions (“simple” endings). We examined the *egl-1* mutant for dendritic tip morphology and found that it had “simple”, unbranched tips more often than wild-type animals (22% simple vs. 6.3% simple, p<0.01 Fisher’s exact test) (Figure 2C). However, the effect size was much smaller relative to what we found in other genetic background and environmental conditions. Moreover, the effect could be due to a number of other factors such as supernumerary cells in the head region of the *egl-1* mutant interfering with proper tip elaboration in certain individuals. Consistent with this idea, we found that *ced-3(n1286)* mutant worms had a similar tip elaboration defect (18% simple), so we did not characterize this phenotype further in the *egl-1* mutant.

Finally, *egl-1* has previously been suggested to have a role in synapse pruning during the development of the RME neurons (MENG *et al.* 2015). Using a GFP-tagged version of the synaptic marker RAB-3 expressed specifically in the URX neurons (BOUNOUTAS *et al.* 2009), we visually compared URX synapses in the nerve ring in wild type and the *egl-1* mutant, but found no gross defects in synapse localization or abundance in the *egl-1* mutant. This suggests that *egl-1* does not play an obvious role in synapse pruning in URX under basal circumstances (Figure 2D).

### *egl-1* expression in URX neurons is activity dependent

Given that previous studies of *egl-1* transcriptional regulation have revealed conserved regulators that are involved in apoptosis in higher organisms, and the novel non-apoptotic nature of the *egl-1* transcription in URX, we set out to characterize how this transcription was controlled. We used the *egl-1* reporter to conduct an unbiased forward genetic screen looking for mutants in which mCherry fluorescence was diminished or lost in URX. We recovered several mutants, including a missense mutation in *cng-1* (CHO *et al.* 2005; WOJTYNIAK *et al.* 2013). The *vx3* allele of *cng-1* is an A-to-C transversion in the third exon, changing an aspartic acid to an alanine. We confirmed the causal nature of this mutation by cell-specific rescue in URX (Figure 3A), and by phenocopy with the canonical *jh111* deletion allele of *cng-1* (data not shown).

**Figure 3.**
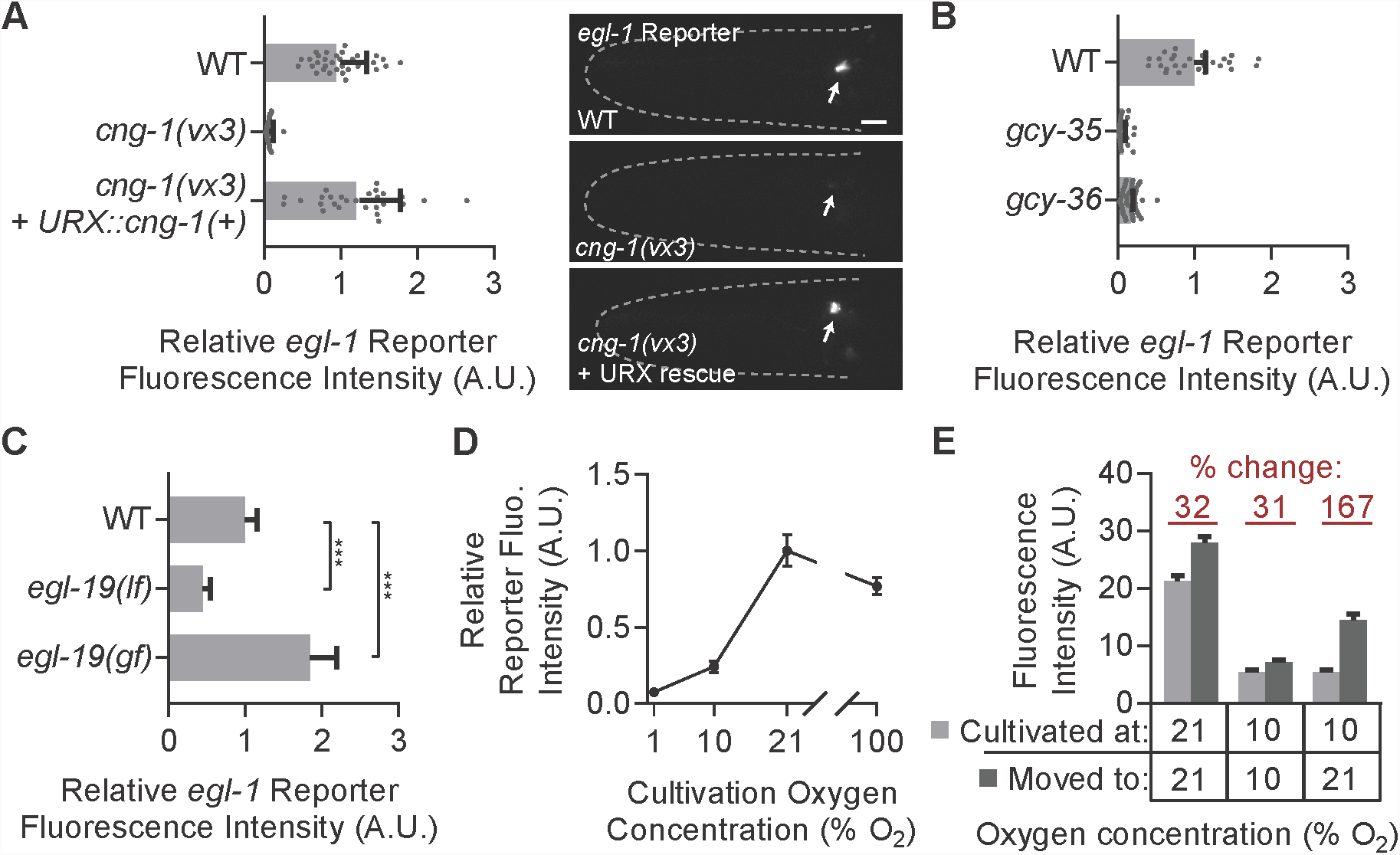
Neuronal activity drives expression of the *egl-1* reporter in the oxygen-sensing URX neurons. ***A.*** *egl-1* reporter expression in day 1 wild-type, *cng-1(vx3)*, and *cng-1(vx3)* worms carrying a *Pgcy-32::cng-1(+)* rescuing transgene and grown at 21% oxygen. Values are normalized to wild type. n ≥ 30 worms per genotype. Representative images are shown to the right; arrows indicate the URX cell body, scale bar is 10 µm. ***B.*** *egl-1* reporter expression in day 1 wild-type worms and mutants lacking components of the oxygen-sensing GCY-35/GCY-36 complex grown at 21% oxygen. Values are normalized to wild-type. n ≥ 30 worms per genotype. ***C.*** *egl-1* reporter expression in day 1 wild-type, *egl-19(n582lf)*, and *egl-19(ad2368gf)* worms. Values are normalized to wild-type. p < 0.01 compared to wild type for both as determined by Student’s t-test, n ≥ 50 worms per genotype. ***D.*** *egl-1* reporter expression in wild-type worms grown until day 1 of adulthood in the indicated oxygen level. Values are normalized to wild-type grown at 21% oxygen. n ≥ 60 worms per condition. **E.** Wild-type populations were reared at the “cultivated at” oxygen level until day 1 of adulthood when *egl-1* reporter expression was quantified. Plates were then moved to the “moved to” oxygen conditions until day 2 of adulthood and *egl-1* reporter expression was again quantified. Bars represent pooled results from three separate trials run on different days, n ≥ 170 worms per condition. All error bars are 95% C.I.

CNG-1 is a cyclic nucleotide-gated ion channel that is an essential component of the oxygen sensation pathway in URX (BUSCH *et al.* 2012; COUTO *et al.* 2013). Thus we hypothesized that perhaps *egl-1* expression was driven by oxygen sensation and neuronal activity. We tested this hypothesis by quantifying reporter expression in other mutants defective in oxygen sensation, as well as in wild-type worms maintained in varying levels of environmental oxygen.

First, we quantified *egl-1* reporter expression in mutants lacking *gcy-35* or *gcy-36*. These genes encode soluble guanylyl-cyclases that heterodimerize and produce cGMP in response to binding molecular oxygen (CHEUNG *et al.* 2004). Both *gcy-35* and *gcy-36* are necessary for URX to become activated in response to changes in environmental oxygen. We found that both *gcy-35* and *gcy-36* mutants had a near complete lack of *egl-1* reporter expression in URX (Figure 3B).

Prolonged calcium entry into URX through CNG-1 in response to long-term oxygen sensation eventually gates the only L-type voltage-gated calcium channel in *C. elegans*, EGL-19 (BUSCH *et al.* 2012). We tested whether EGL-19 was necessary for *egl-1* expression in URX by quantifying *egl-1* reporter expression in both a loss-of-function allele of *egl-19*, as well as a gain-of-function allele which has an increased open probability compared to native EGL-19 (LAINE *et al.* 2014). We found that *egl-1* reporter expression in URX was significantly decreased in the *egl-19(n582)* loss-of-function allele, and significantly increased in the *egl-19(n2368)* gain-of-function allele, signifying that EGL-19 activation downstream of prolonged oxygen sensation is necessary for *egl-1* transcription in URX (Figure 3C).

We next directly assessed whether environmental oxygen sensation drives *egl-1* expression in URX by cultivating wild-type animals with the *egl-1* reporter in different oxygen environments and quantifying reporter expression on day 1 of adulthood. Wild isolates of *C. elegans* have a preference for 5-12% O2 conditions, which is thought to reflect the environments most favorable to their survival (GRAY *et al.* 2004). These strains will avoid both hypoxia (<1% O2) and oxygen levels over 12% O2, including the 21% O2 conditions normally found at the lab benchtop. The lab strain N2 has a dampened capability to avoid 21% oxygen due to a gain-of-function mutation in the gene *npr-1* (DE BONO AND BARGMANN 1998). This difference in behavior is not explained by differences in the URX neuron itself however, as both N2 and the *npr-1* loss-of-function mutant (which mimics the *npr-1* allele found in natural isolates) display similar URX calcium dynamics in response to oxygen (JANG *et al.* 2017). Instead, the difference in behavior is thought to originate due to functional differences downstream of URX. For these reasons, we grew animals in 1%, 10%, or 21% oxygen environments. We also grew worms in 100% oxygen to test whether the *egl-1* reporter would be expressed higher than at 21% conditions. This oxygen condition is likely much higher than worms would experience in their natural environment.

We found that *egl-1* reporter expression levels reflected the cultivation level of oxygen up to 21% oxygen, with day 1 animals grown in 1% oxygen having 0.08% the level of reporter expression as those grown in 21% oxygen (Figure 3D). Interestingly, worms grown at 100% oxygen actually had slightly lower expression of the reporter than those grown at 21%, and the morphology of URX was normal in these worms.

We considered the possibility that growth from egg in these different environments had caused developmental differences in the URX neuron that led to the differences we observed in reporter expression, so we carried out a complementary set of experiments where populations were grown at one oxygen concentration until day 1 of adulthood and then shifted to another oxygen level until day 2. Worms grown at 10% or 21% oxygen, and then maintained at those same levels until day 2 had an increase of reporter expression of about 30% over that period, while populations grown at 10% oxygen then moved to 21% oxygen had a mean increase of 167% from day 1 to day 2 (Figure 3E). Taken together, these experiments strongly suggest that environmental oxygen sensation by URX is necessary for expression of *egl-1* in URX.

### EGL-4 and KIN-29 regulate *egl-1* expression in URX partially in parallel

We undertook two approaches to identify other genetic pathways that may regulate *egl-1* expression in URX: a reversion screen where we screened for mutants that recovered *egl-1* reporter expression in a *cng-1(vx3)* mutant background, and a candidate screen focused on other genes known to regulate activity-dependent genes.

Our reversion screen recovered several alleles of *egl-4*, the worm homolog of Protein Kinase G, as determined by complementation tests and phenocopy (DANIELS *et al.* 2000; STANSBERRY *et al.* 2001; L’ETOILE *et al.* 2002). Both *egl-4(vx19);cng-1(vx3)* double and *egl-4(n477)* single mutants expressed the *egl-1* reporter higher than wild-type (Figure 4A, 4D). This indicates that EGL-4 likely acts downstream of CNG-1 to repress expression of *egl-1*. This is consistent with past studies that have outlined how EGL-4 translocates into the nucleus to affect gene expression in response to long-term neuronal activity (O’HALLORAN *et al.* 2012). We confirmed expression of *egl-1* in URX in the *egl-4(n477)* mutant by smFISH (Figure 4B), further affirming the fidelity of the *egl-1* reporter to endogenous *egl-1* transcription.

**Figure 4.**
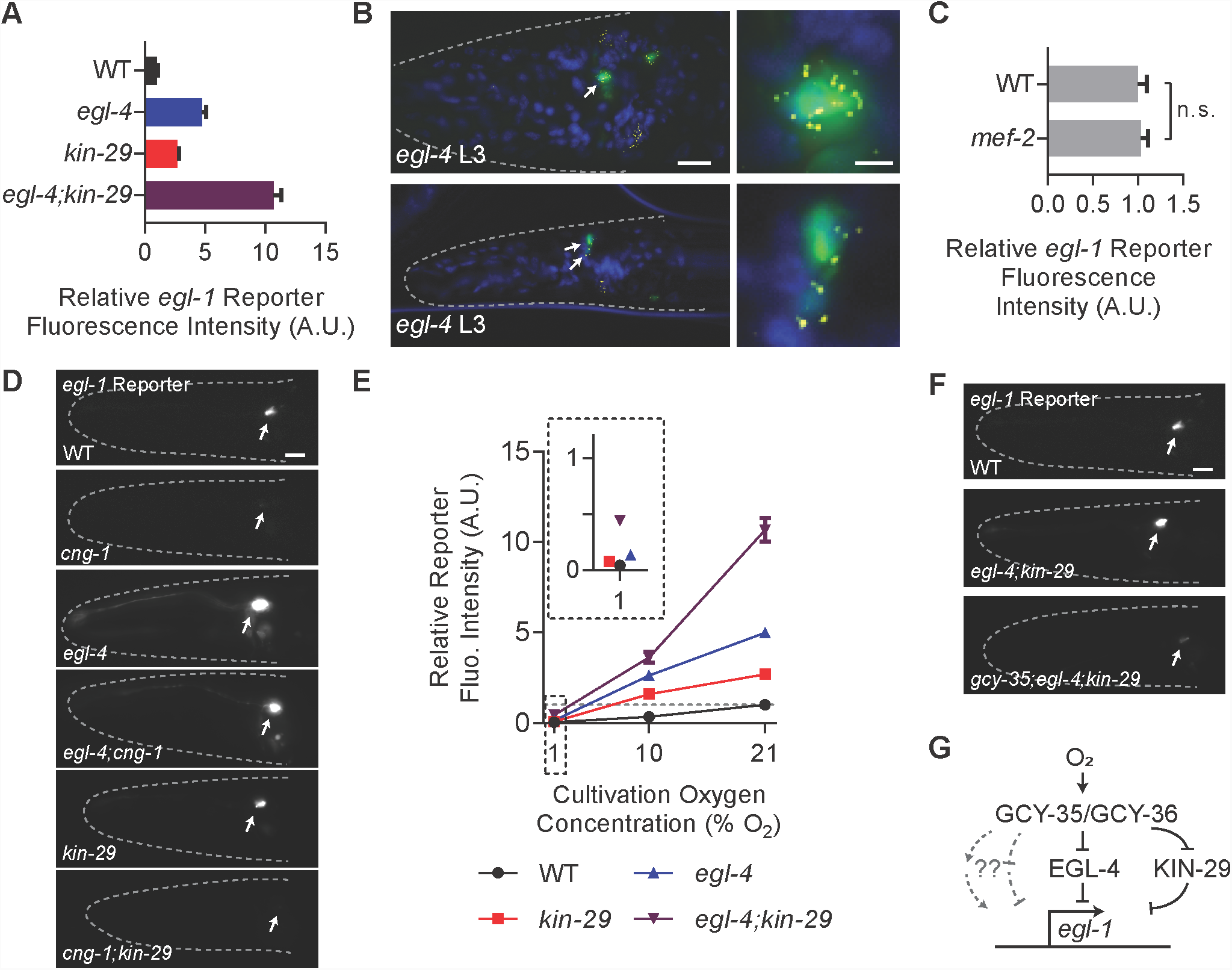
At least three pathways regulate *egl-1* expression in URX neurons. ***A.*** *egl-1* reporter expression at 21% oxygen in animals of the given genotype normalized to WT. Animals were synchronized by timed egg-laying and then imaged 3 days later. n ≥ 55 worms per genotype. **B.** smFISH showing *egl-1* transcripts in *egl-4(n477)* mutant worms carrying the *muIs102* reporter (*Pgcy-32::GFP*) to label URX green. Blue is DAPI, *egl-1* mRNA transcripts are labeled yellow. Genotype and larval age of each animal is shown in white text. ***C.*** *egl-1* reporter expression in day 1 wild type and *mef-2(gv1)* worms at 21% oxygen, values normalized to wild type. n ≥ 45 worms per genotype. **D.** Representative images of day 1 adult WT N2, *cng-1(vx3), egl-4(n477), egl-4(vx19);cng-1(vx3), kin-29(oy38)*, and *cng-1(jh111);kin-29(oy38)* worms. *cng-1(vx3)* and *cng-1(jh111)* alleles fail to complement, as do *egl-4(n477)* and *egl-4(vx19)*. ***E.*** *egl-1* reporter expression in the given genotypes at the given oxygen levels. Animals were synchronized by timed egg-laying and then imaged 3 days later. Values are normalized to a yoked wild-type plate grown at 21% oxygen. n ≥ 55 worms per genotype per condition. **F.** Representative images of day 1 adults of the given genotypes. **G.** Model showing a possible configuration of the relationship between oxygen sensation in URX and *egl-1* transcription. Arrows and lines do not imply direct interaction, only the valence and order of the genetic pathway. The grey dotted lines with question marks represent two possibilities for the third, oxygen-dependent pathway our epistasis analysis suggests. All error bars are 95% C.I. In photos, arrows indicate URX cell body and scale bars are 10 µm, except for the enlargements in panel B, in which the scale bars is 2µm.

Separately, we also examined *egl-1* reporter expression in another kinase pathway mutant that has been previously implicated in regulating activity-dependent gene expression. KIN-29 is the worm homolog of the <u>S</u>alt-Inducible <u>K</u>inase (SIK) family of proteins, and like EGL-4, has been shown to translocate to the nucleus to affect gene transcription in certain contexts (KATOH *et al.* 2002; VAN DER LINDEN *et al.* 2007; VAN DER LINDEN *et al.* 2008). We found that the *kin-29* mutant had increased *egl-1* reporter expression in URX compared to wild-type, though less than the *egl-4* mutant (Figure 4A).

EGL-4 and KIN-29 have previously been shown to regulate expression of the *str-1* chemoreceptor in the AWB sensory neuron by phosphorylating the MEF-2 transcription factor, antagonizing its activity. If MEF-2 were also involved in expression of *egl-1* in URX, we would expect it to have the opposite phenotype of EGL-4, which would be decreased expression of the *egl-1* reporter at 21% oxygen. However, the *mef-2(gv1)* mutant had the same level of *egl-1* reporter expression as wild type (Figure 4C). This suggests that EGL-4 likely does not act through MEF-2 in regulating *egl-1* expression as it does *str-1*.

Several pieces of evidence suggest that *egl-4* and *kin-29* act at least partially in parallel to regulate *egl-1* transcription in URX. First, we found that the *egl-4;kin-29* double mutant had higher *egl-1* reporter expression than either single mutant (Figure 4A). Reporter intensity at 21% oxygen normalized to wild type was 10.7 ± 2.6 (st. dev.) for the *egl-4;kin-29* double mutant, compared to 5.0 ± 1.0 for *egl-4* and 2.7 ± 0.6 for *kin-29* single mutants. Also, we found that the *cng-1;kin-29* double mutant had very low expression of the *egl-1* reporter, in contrast to the *egl-4;cng-1* double mutant which had high expression of the reporter (Figure 4D). These results strongly suggest that *egl-4* and *kin-29* act through partially separate pathways to repress expression of *egl-1* in URX.

Above, we found that the *egl-1* reporter was expressed at a high level in the *egl-4* mutant even when the URX oxygen transduction pathway was severely compromised with the *cng-1* mutation. This made us wonder whether environmental oxygen could still regulate *egl-1* transcription in URX in mutants without the *egl-4* and *kin-29* repressor pathways. We hypothesized that the *egl-1* reporter would be expressed highly even at low oxygen in these mutants. However we found that *egl-4* and *kin-29* single mutants as well as the *egl-4;kin-29* double mutant still showed decreased reporter expression in low oxygen conditions (Figure 4E). This provides evidence for at least a third, oxygen-dependent mechanism to regulate *egl-1* expression in URX. As an independent test for this hypothesis, we asked if the *egl-1* reporter was still expressed in *egl-4;kin-29* after mutation of the oxygen sensor GCY-35. We found that the triple mutant *gcy-35;egl-4;kin-29* showed vastly decreased reporter expression compared to the *egl-4;kin-29* double mutant (Figure 4F). These results are consistent with two possibilities: either oxygen sensation by URX is necessary to drive *egl-1* expression or there is at least one repressive pathway other than the *egl-4* and *kin-29* pathways that can regulate *egl-1* expression in URX in an oxygen-dependent manner (Figure 4G).

## DISCUSSION

### *egl-1* is transcribed in the URX neurons post-developmentally without killing them

Here we described the transcription of the proapoptotic BH3-only gene *egl-1* in the URX neuron pair in post-developmental *C. elegans*. This expression is unusual because in stands in contrast to the typical context of *egl-1* expression, which mostly occurs during development to trigger apoptosis. Previous work has suggested that both pro- and anti-apoptotic genes are present in developing cells (SHAHAM AND HORVITZ 1996). They function antagonistically, keeping a cell alive unless the apoptotic cascade is initiated by *egl-1* expression. However, *egl-1* transcription in URX, which continues past when somatic cell deaths in the hermaphrodite worm cease, does not lead to cell death in any of the circumstances we examined.

In mammals, several BH3-only proteins are expressed in the adult animal in various tissues, including the nervous system (O’REILLY *et al.* 2000; NAPANKANGAS *et al.* 2003; COULTAS *et al.* 2004; STURM *et al.* 2006; LEIN *et al.* 2007; HAWRYLYCZ *et al.* 2012). It is difficult to assess the implication of this expression because many mammalian BH3-only proteins are regulated post-transcriptionally; however, this area of study is largely unexamined and could prove to be of significance. With at least eight family members, studying BH3-only proteins in mammals can be difficult due to redundancy in their functions (GIAM *et al.* 2008). In contrast, *C. elegans* has only two BH3-only proteins, *egl-1* and *ced-13* (SCHUMACHER *et al.* 2005). EGL-1 is necessary for nearly all developmental somatic cell deaths while CED-13 seems to function in germline cell death and does not appear to overlap with the activity of *egl-1* (SCHUMACHER *et al.* 2005; KING *et al.* 2018). These factors make studying the roles of BH3-only proteins more straightforward in *C. elegans*.

Intriguingly, transgenic expression of *egl-1* has been previously used to genetically ablate the URX neurons. The transgene array *qaIs2241* drives *egl-1* expression from the *gcy-36* promoter, and the death of URX in these animals was evident by both visual inspection and by URX-specific behavioral deficits (CHANG *et al.* 2006; STYER *et al.* 2008; CARRILLO *et al.* 2013; ZHAO *et al.* 2018). Albeit artificial, this shows that URX is capable of undergoing apoptosis in an *egl-1*-dependent manner.

We propose two possibilities to explain why this *egl-1* transgene array kills URX while the endogenous expression of *egl-1* we describe in this study does not. The first possibility relates to the expression level of *egl-1* in URX. The *gcy-36* gene is highly expressed in URX and *qaIs2241* contains multiple copies of *Pgcy-36::egl-1* in an array, so transgenic *egl-1* is likely expressed much higher than what occurs endogenously. The second possibility is more speculative and relates to expression timing. Because we found that endogenous *egl-1* expression depends on GCY-36, it is likely that endogenous *egl-1* expression must necessarily begin after *gcy-36* expression. By contrast, transgenic *egl-1* expressed from *qaIs2241* under the control of the *gcy-36* promoter loses this dependence, expressing earlier than the endogenous *egl-1*. Note that these two possibilities are not mutually exclusive.

### *egl-1* transcription in URX is regulated by neuronal activity, EGL-4, and/or KIN-29

We found that sensory transduction, the worm PKG homolog EGL-4, and the worm SIK homolog KIN-29 all regulate *egl-1* expression in URX in a complex manner. The cellular localization of EGL-4 has previously been studied directly in AWC sensory neurons, where it was shown to translocate to the nucleus when intracellular cGMP decreased with prolonged exposure to an AWC-sensed odorant (O’HALLORAN *et al.* 2012). Interestingly, URX has a constitutively high level of cGMP in wild-type animals at 21% oxygen and even higher cGMP production in a *cng-1* mutant background (COUTO *et al.* 2013), yet our epistasis analysis found that EGL-4 repressed *egl-1* expression in URX when *cng-1* was mutated. This suggests that EGL-4 localization may be regulated differently in URX than in AWC, and that calcium entry may play a role. EGL-4 and KIN-29 have also been found to work together to regulate other activity-dependent genes, including several chemoreceptors in subsets of *C. elegans* neurons (VAN DER LINDEN *et al.* 2007; VAN DER LINDEN *et al.* 2008). We found that, unlike how they are proposed to work in AWB to regulate expression of the *str-1* chemoreceptor, EGL-4 and KIN-29 likely do not act through the MEF-2 transcription factor to regulate *egl-1* expression. Also, the valence of expression for *egl-1* is opposite that of *str-1* in *egl-4* and *kin-29* mutants. That is, while *str-1* is downregulated in *egl-4* and *kin-29* mutants, *egl-1* is upregulated in these same genetic backgrounds. These differences with another known activity-dependent gene governed by *egl-4* and *kin-29* suggest that studying *egl-1* expression in URX can offer a complementary system to identify factors that regulate activity-dependent gene expression in *C. elegans*.

Both Protein Kinase G and the Salt-Inducible Kinase family have been shown to be involved in cell death regulation, in both repressive and activating roles in other model systems (FISCUS 2002; DEGUCHI *et al.* 2004; CHENG *et al.* 2009; FALLAHIAN *et al.* 2011; DU *et al.* 2016; TARUMOTO *et al.* 2018) – however, their relation to BH3-only proteins has largely not been investigated. Our results suggest that exploring whether PKG and SIK family members regulate BH3-only proteins in other systems may be a worthwhile area of study. Though the transcription of *egl-1* in URX that we describe does not lead to apoptosis, it is feasible that conserved regulators in other animals may have evolved a general or circumstantial role in programmed cell death.

### The *egl-1* mutant is not impaired for multiple URX-related phenotypes

One important question that remains unanswered is whether or not the *egl-1* mRNA we see in URX is translated to a protein. We attempted to answer this question using a variety of approaches, such as immunohistochemistry using commercially available antibodies, tagging the endogenous protein with CRISPR, and transgenically expressing a tagged version of the protein. Unfortunately these attempts were unsuccessful, so we were unable to definitively assess whether the *egl-1* protein product is present in URX. Other labs have also reported being unable to tag the EGL-1 protein (B. Conradt, personal communication).

Despite the lack of a clear answer about the presence of EGL-1 protein in URX, we nevertheless compared the *egl-1* mutant and wild-type worms with regard to the known URX-related behaviors of bordering due to 21% oxygen sensation and immune response to pathogenic bacteria. For both cases, we found no differences. We also examined URX at the cellular level in the *egl-1* mutant by looking at dendrite and synapse morphology. General synapse morphology looked normal in the *egl-1* mutant, though there was a slight defect in dendritic tip morphology compared to wild type. However, this defect could have several origins that are unrelated to an action of *egl-1* in URX, so we did not follow this subtle phenotype further.

In the absence of a clear abnormal phenotype of the *egl-1* mutant with regard to URX function, we can only speculate about why *egl-1* is transcribed in this living pair of neurons. One possibility is that whereas *egl-1* normally triggers cell death during development, its expression in URX represents an atypically regulated form of apoptosis for which the *egl-1* transcript is maintained in the cell so that in response to some additional trigger the translated protein product then induces apoptosis. There is at least one other example for which *egl-1* expression does not cause immediate cell death (JOHNSEN AND HORVITZ 2016); the cell B.alrapaav in the male tail expresses *egl-1* but persists in a poised state unless a neighboring cell begins to engulf it, at which point cell death occurs – a process termed “assisted suicide.” The URX neuron pair might similarly exist in a poised state in the wild-type worm. In this model, one or more unknown signals would act as the “trigger” that causes translation of EGL-1 and possibly cell death. Because wild isolates of *C. elegans* typically prefer 5-12% O2 conditions and URX is tonically active, one trigger might be prolonged exposure to higher concentrations of oxygen. However, URX neurons show no evidence of death or dysfunction when cultivated at 21% or 100% oxygen. Also, we found that exposure to 100% oxygen did not lead to higher *egl-1* reporter expression than at 21% oxygen. Thus, high oxygen levels do not appear to serve as an apoptotic trigger for URX, at least alone.

One potential problem with this apoptotic trigger hypothesis, however, is that the canonical downstream binding partner of EGL-1, CED-9, and the downstream effector CED-4 are both thought to no longer be expressed in somatic tissue past development (CHEN *et al.* 2000), so EGL-1 protein expressed in URX in late larval and adult *C. elegans* might not induce apoptosis.

An alternate possibility is that *egl-1* expression in URX represents a moonlighting function for EGL-1, similar to the non-apoptotic roles that are beginning to be uncovered for mammalian BH3-only proteins (ESPOSTI *et al.* 2001; DANIAL *et al.* 2003; MAIURI *et al.* 2007; DANIAL *et al.* 2008; SINHA *et al.* 2008; GIMENEZ-CASSINA *et al.* 2014; LINDQVIST *et al.* 2014). The apoptotic-trigger and moonlighting possibilities are not mutually exclusive.

Activity-dependent *egl-1* transcription in post-developmental *C. elegans* might provide a new system for the study of mechanisms that regulate activity-dependent gene expression and might also possibly reveal a non-apoptotic role for a BH3-only protein. Our future work will focus on identifying the mechanisms and transcription factors by which neuronal activity controls *egl-1* transcription in URX, and on what role *egl-1* might play in this neuron.

## ACKNOWLEDGMENTS

We would like to thank Lina Gomez and Luisa Scott for helpful discussions, Michael Nonet for reagents, and Susan Rozmiarek for expert assistance. Some strains were provided by the CGC, which is funded by NIH Office of Research Infrastructure Programs (P40 OD010440). Additional funds were provided by NIH-NIA grants RF1AG057355 and R01AG041135. V.K.D. was a Howard Hughes Medical Institute International Student Research fellow. H.R.H. is an Investigator of the Howard Hughes Medical Institute.

---

**Figure S1.**
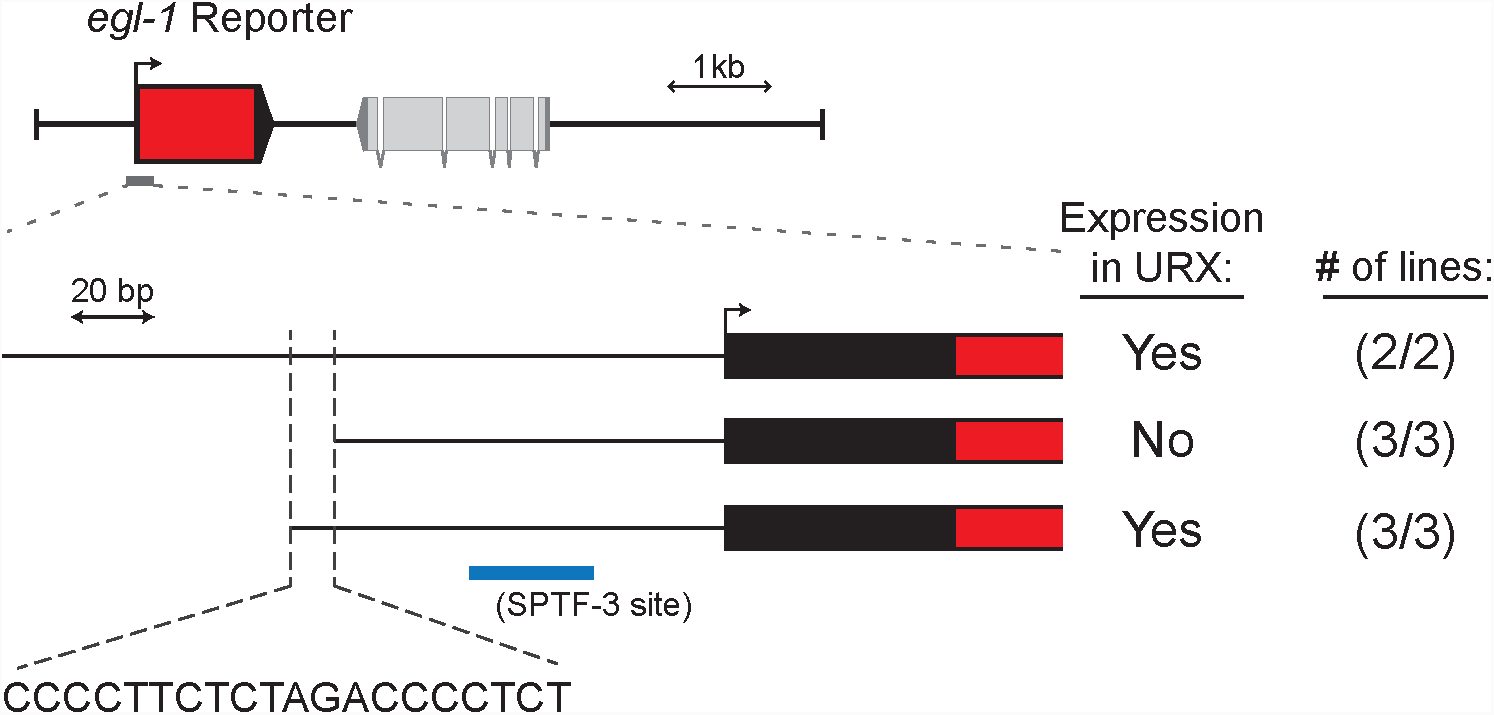
Promoter truncations to identify *egl-1* regulatory region necessary for URX expression. Top shows a schematic of the *egl-1* reporter after replacement of the *egl-1* genomic region with *mCherry*. Grey line indicates area enlarged below. The *egl-1* 5’UTR is shown in black, *mCherry* is shown in red, and the transcriptional start site is shown with a bent arrow.

